# A role for the F-box protein HAWAIIAN SKIRT in plant miRNA function

**DOI:** 10.1101/123703

**Authors:** Patricia L.M. Lang, Michael D. Christie, Ezgi Dogan, Rebecca Schwab, Jörg Hagmann, Anna-Lena Van de Weyer, Detlef Weigel

## Abstract

As regulators of gene expression in multicellular organisms, microRNAs (miRNAs) are crucial for growth and development. While a plethora of factors involved in their biogenesis and action in *Arabidopsis thaliana* have been described, these processes and their fine-tuning are not fully understood in plants. Here, we used plants expressing an artificial miRNA target mimic (MIM) to screen for negative regulators of miR156 activity. We identified a new mutant allele of the F-box protein HAWAIIAN SKIRT (HWS; At3G61590), *hws-5*, as a suppressor of the *MIM156*-induced developmental and molecular phenotypes. In *hws* plants, levels of endogenous miRNAs are increased and their mRNA targets decreased. Plants constitutively expressing full-length HWS - but not a truncated version lacking the F-box domain - display morphological and molecular phenotypes resembling those of mutants defective in miRNA biogenesis and activity. In combination with such mutants, *hws* loses its delayed floral organ abscission (‘skirt’) phenotype, suggesting epistasis. Also, the overall *hws* transcriptome profile partially resembles well-known miRNA mutants *hyl1*-2 and *se-3*, indicating action in a common pathway. We thus propose HWS as a novel, F-box dependent regulator of miRNA biogenesis.

**Summary statement:** HAWAIIAN SKIRT is a regulator of *Arabidopsis thaliana* microRNA biogenesis and acts in an F-box-dependent manner.

## Introduction

MicroRNAs (miRNAs) are 21 to 24 nucleotide (nt) long single-stranded RNA molecules that are crucial for regulation and fine-tuning of gene expression in multicellular organisms (Bartel 2009). In plants, transcription of *MIRNA* genes generates longer precursor RNAs with characteristic stem-loop folding, and is followed by two major nuclear processing (‘dicing’) steps mediated mainly by the endoribonuclease DICER-LIKE1 (DCL1), and aided by cofactors including SERRATE (SE) and HYPONASTIC LEAVES1 (HYL1) (Kurihara & Watanabe 2004; Laubinger et al. 2008; Vazquez et al. 2004; Bernstein et al. 2001). The first dicing step, usually near the base of the precursor stem, excises the primary miRNA stem-loop, which is cut once more to remove the loop and form the duplex of miRNA and miRNA* strand. After 3’-O methylation and translocation to the cytoplasm (Yu et al. 2005), the mature miRNA, or guide, strand associates with an ARGONAUTE (AGO) protein and together they form an active miRNA INDUCED SILENCING COMPLEX (miRISC). By sequence complementarity of the miRNA to its mRNA targets, miRISC selectively inhibits translation or triggers transcript cleavage (Rogers & Chen 2013; Brodersen et al. 2012).

Several mechanisms that control the levels of miRNAs and thereby adjust target silencing have been described: transcriptional regulation ensures the correct spatio-temporal expression of *MIR* genes, while post-transcriptional steps further fine-tune miRNA accumulation (Achkar et al. 2016). Additionally, mature miRISC activity can be attenuated by miRNA target mimicry (Franco-Zorrilla et al. 2007). There, a non-coding RNA containing a sequence motif that is complementary to a miRNA sequesters the respective miRISC and thus renders it unavailable for inhibition of regular mRNA targets (Franco-Zorrilla et al. 2007). In animals, this principle is known as competing endogenous RNAs (ceRNAs), although its exact significance as regulatory mechanism appears to remain controversial (Wang et al. 2015; Bak & Mikkelsen 2014; Denzler et al. 2016). Only a single case of natural miRNA target mimicry, that of *IPS1*, which interferes with the activity of miR399, has been studied in detail in plants. However, positive posttranscriptional regulation by mismatched miRNAs associating with their target and thereby protecting it, has been proposed as a mimicry related mechanism (Couzigou et al. 2016).

The miRNA target mimicry principle has been extensively used to investigate the biological function of miRNAs (Franco-Zorrilla et al. 2007; Sha et al. 2014). Artificial miRNA target mimicry lines (MIMs) based on the endogenous *INDUCED BY PHOSPHATE STARVATION1* (*IPS1*) transcript, or on entirely artificial sequences, have been used to interfere with the majority of *Arabidopsis thaliana* miRNAs (Franco-Zorrilla et al. 2007; Todesco et al. 2010; Yan et al. 2012; Reichel et al. 2015). Many conserved plant miRNAs are encoded by medium-size gene families (Li & Mao 2007). In *A. thaliana*, one such family is the deeply conserved miR156 family with eight *MIR* genes, encoding miR156a to miR156h (Griffiths-Jones 2004; Kozomara & Griffiths-Jones 2014). Mature miR156 accumulates in the shoots of juvenile plants and gradually decreases in abundance as the plant ages (Wang 2014; Wang et al. 2009; Wu et al. 2009; Wu & Poethig 2006). It directly regulates at least 10 out of 16 members of the SQUAMOSA PROMOTER-BINDING PROTEIN-LIKE (SPL) transcription factor family, which control a range of biological processes, most of them relating to developmental progression during the vegetative phase of plant growth (Xu et al. 2016).

Plants constitutively expressing the *IPS1*-based *MIM156* show reduced miR156 activity and display an accelerated juvenile-to-adult phase transition as well as exaggerated adult growth traits such as serrated leaf margins (Franco-Zorrilla et al. 2007). Similar phenotypes are observed in plants expressing miR156-insensitive versions of SPL targets (Wang et al. 2008; Wu & Poethig 2006; Franco-Zorrilla et al. 2007). In those plants, the rate at which rosette leaves are initiated during vegetative growth is also greatly reduced, and *MIM156* cotyledons are bent and spoon-shaped (Todesco et al. 2010). In contrast, ectopic overexpression of miR156, which reduces SPL levels, prolongs the juvenile phase with non-serrated rosette leaves, and plants initiate rosette leaves faster than the wild-type (Wu & Poethig 2006).

We sought to identify negative regulators of miR156 activity by screening for suppressors of *MIM156*-induced developmental phenotypes. Here, we describe the F-box protein encoded by *HAWAIIAN SKIRT* (*HWS*, At3G61590) as a broad suppressor of miRNA activity in *Arabidopsis thaliana*. As part of an Skp-Cullin-F-box (SCF) complex, F-box proteins provide E3 ubiquitin ligases with target specificity via recognition of substrates for ubiquitination (Risseeuw et al. 2003). HWS was previously shown to interact with the classical components of an SCF complex in a yeast-two-hybrid (Y2H) screen. Those are ASK20A and ASK20B, the two translational products of *ARABIDOPSIS SKP1-like 20* (*ASK20*), and ARABIDOPSIS SKP-LIKE1 (ASK1), which functioned as bridge between CULLIN1 (CUL1) and HWS (Ogura et al. 2008; Kuroda et al. 2002). To date, no substrates for HWS, marked either for proteasome-mediated degradation or other molecular processes, have been identified. Mutations in *HWS* affect root growth via regulation of quiescent-center independent meristem activity as well as stomatal guard cell development (Kim et al. 2016; Yu et al. 2015). Due to delayed abscission and sepal fusion, *hws* mutants also fail to shed sepals, petals and anthers (Gonzalez-Carranza et al. 2007). Loss of *HWS* furthermore results in increased organ growth, whereas overexpression yields smaller plants with elongated, serrated and hyponastic leaves, thus resembling phenotypes previously described for the miRNA biogenesis mutants *hyl1-2, dcl1-100* or *hst1* as well as hypomorphic *ago1* alleles (Gonzalez-Carranza et al. 2007).

Our study demonstrates that *HWS* also broadly affects plant miRNA function. Mutations in *HWS* increase the steady-state levels of several different miRNAs, resulting in corresponding decreases in the levels of their respective targets. Overexpression of full-length HWS has an inverse effect on the levels of miRNAs and their targets, and this function requires its F-box domain. The characteristic delayed floral organ abscission, or ‘skirt’ phenotype of *hws* mutants is lost when combined with mutants defective in miRNA biogenesis. This indicates that *HWS* and miRNA factors like *SE, HYL1* and *AGO1* are epistatic to each other and active in a common pathway, a notion supported by significant overlaps within *hws* and miRNA mutant transcript profiles. We propose that HWS is a new factor involved in the biogenesis of miRNAs, and exerts its function through an F-box dependent process.

## Results

### *hws* mutations suppress miRNA target mimicry-induced phenotypes

To identify genetic modifiers that suppress the effects of an *IPS1-based MIM156* transgene, we focused on three easily monitored developmental abnormalities characteristic of lines ectopically expressing this transgene: spoon-shaped cotyledons, premature rosette leaf serration, and a reduced leaf initiation rate during vegetative growth. One line that we found after EMS mutagenesis displayed broad suppression of all *MIM156*-induced developmental abnormalities (Fig. 1A). Using mapping by sequencing, we localized the causal mutation to a region on the right arm of chromosome 3 (Fig. 1A and B; see Materials and Methods for details).

**Figure 1.**
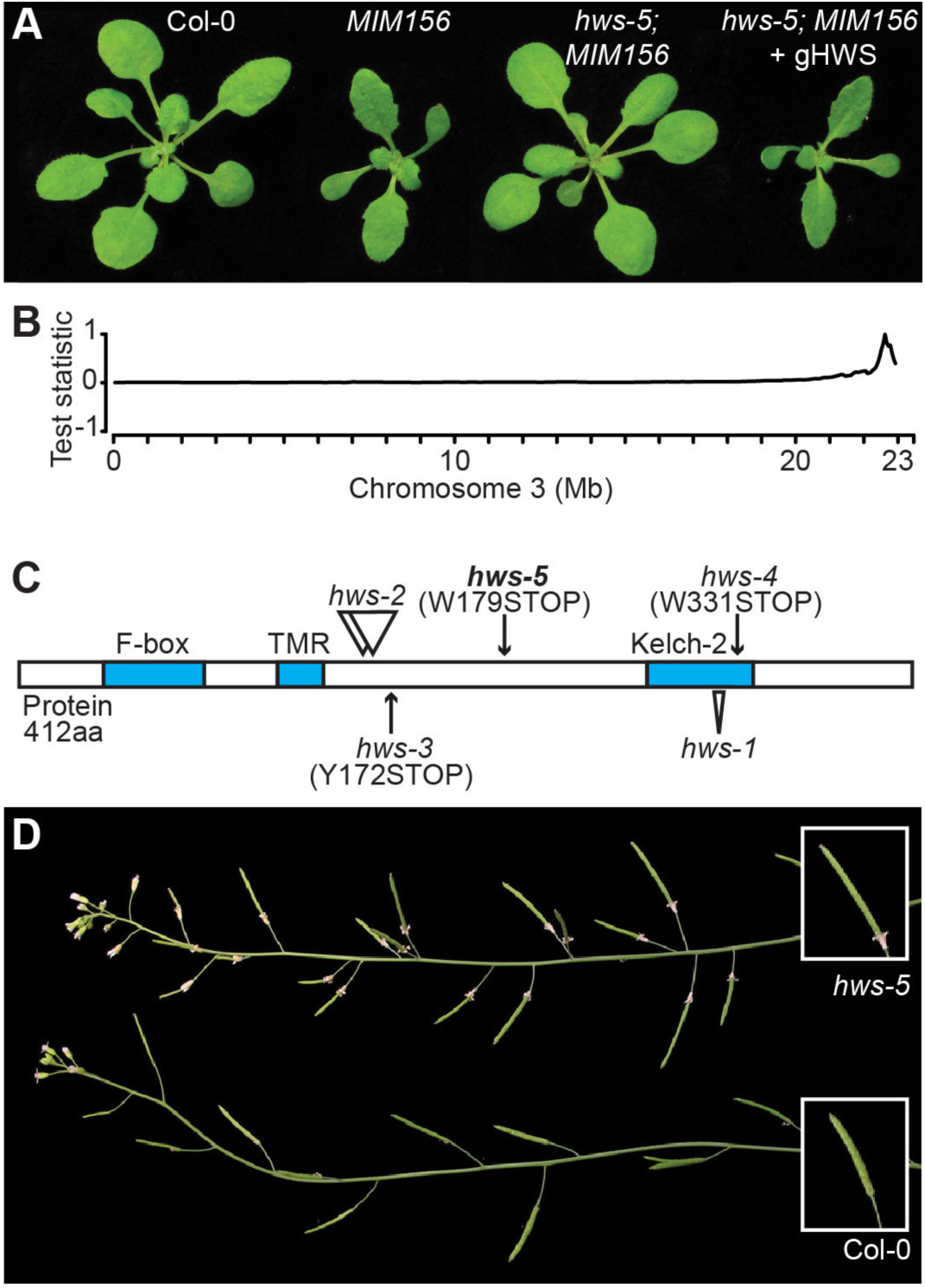
Characterization of the *hws-5* mutant. (*A*) Phenotype of 21-day old Col-0, *MIM156, hws-5; MIM156* and *hws-5; MIM156 + gHWS* plants. (*B*) Chromosome 3 SHOREmap results for *hws-5*. (*C*) Location and effects of the mutations on the HWS protein. Annotated or predicted domains are marked in blue. (*D*) Sepal-fusion “skirt” phenotype and phyllotactic distortion in *hws-5* compared to Col-0.

Within this region, a G-to-A single nucleotide substitution (G537A, chr3:22793585, TAIR10) was identified that caused a premature termination codon (W179STOP) in *HAWAIIAN SKIRT* (*HWS;* At3g61590). HWS is a 412 amino acid (aa) protein with an N-terminal F-box domain, a predicted transmembrane domain, and a C-terminal Kelch-2 domain (Fig. 1C, (Gonzalez-Carranza et al. 2007)). The typical *MIM156* phenotype was restored in mutants transformed with a genomic construct of *HWS*, confirming that *HWS* was indeed the causal locus (Fig. 1A). We henceforth refer to the mutant allele as *hws-5*. Levels of *HWS* transcript detected by reverse transcription quantitative PCR (RT-qPCR) are similarly decreased in *hws-5* as in *hws-1* plants as compared to wild type (see below for description of mutant), suggesting that *hws-5* is a hypomorphic allele (Fig. S1F).

Other *hws* mutant alleles have previously been isolated and shown to affect root-meristem activity (*hws-3* and *hws-4*, (Kim et al. 2016)), and to be impaired in the abscission of floral organs (*hws-1* and *hws-2* (Gonzalez-Carranza et al. 2007)). Incomplete separation of sepals imposes a structural barrier that prevents the shedding of sepals, petals and stamens (Gonzalez-Carranza et al. 2007). Both with and without the MIM156 transgene, we observed similarly impaired abscission in *hws-5* plants (Fig. 1D, Fig. S1A, B and C). Partial fusion of cauline leaves to the inflorescence stem was evident in both *hws-1* and *hws-5* mutants, and like sepal abscission persisted in the presence of the MIM156 transgene (Fig. S1E).

Analyzing total RNA profiles by sequencing (RNA-seq), we identified differentially expressed (DE) genes in the suppressor line *hws-5; MIM156* compared to the isogenic *MIM156* parent (false discovery rate adjusted p-value <0.05). Upregulation of *IPS1* (At3G09922) transcripts in both lines, though more strongly in *MIM156*, confirmed that suppression was not simply due to loss of expression of the *IPS1*-based *MIM156* transgene. Genes differentially expressed between *hws-5; MIM156* versus *MIM156* alone are similar in number compared to those found between *MIM156* and the Col-0 wild type (12% versus 15%, respectively; Fig. 2A). In addition, they are also largely overlapping (76 and 57% of all DE genes respectively) and similarly up- or downregulated in Col-0 and *hws-5; MIM156* when compared to *MIM156* plants (99% of genes in the overlap). These observations are in line with the restoration of a wild-type appearance of *hws-5; MIM156* seedlings. This indicates that the suppression of the *MIM156* morphological phenotype extends broadly to the transcriptional level (Fig. S1G, Table S1).

**Figure 2.**
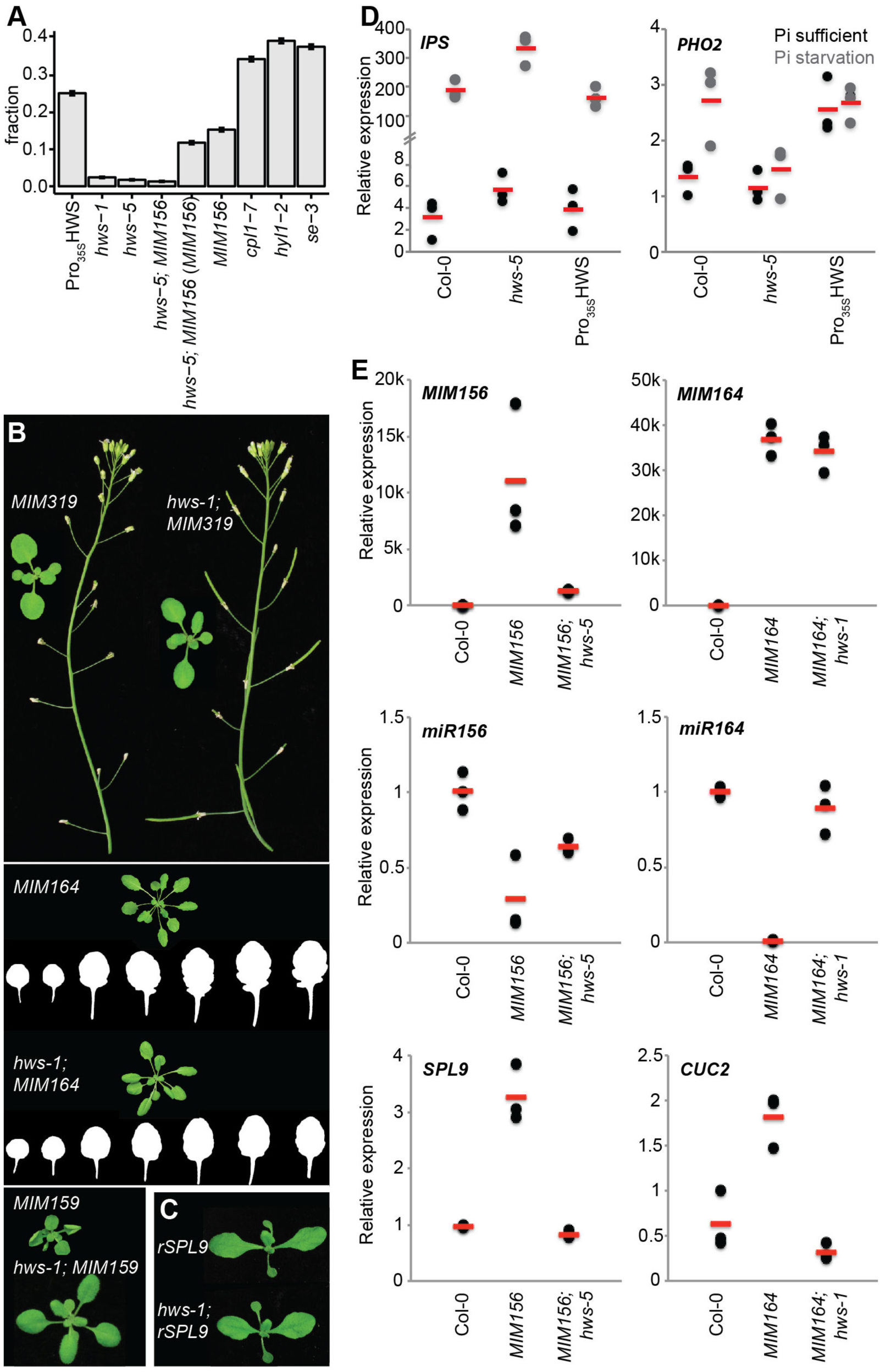
Effects of *hws* mutation on *MIM* transgene-dependent and -independent phenotypes. (*A*) Fraction of differentially expressed genes in *35S::HWS, hws-1, hws-5, hws-5; MIM156* (compared to Col-0 and to *MIM156*), *MIM156* (n=2) and miRNA-mutants *cpl1-7, hyl1-2* and *se-3* (n=3, data from (Manavella et al. 2012)) with 95% confidence intervals. For p-values, see Table S4. (*B*) Suppression of *MIM159, MIM319* and *MIM164* phenotypes in *hws-1* background. (*C*) miR156 resistant *rSPL9* and *rSPL9; hws-5* plants. (*D*) Relative expression of *IPS* and *PHO2* in Col-0, *hws-5* and *hws-1* plants harboring *35S::HWS*. (*E*) Relative expression of *MIM156, miR156, SPL9* in Col-0, *MIM156* and *hws-5; MIM156* and of *MIM164, miR164* and *CUC2* in Col-0, *MIM164* and *hws-1*; *MIM164*. Dots represent biological replicates as means of technical replicates (n=3), bars indicate means of biological replicates.

To determine whether HWS’ activity was specific to miR156, we tested if other mimicry lines were equally affected. We combined *hws-1* with three additional, ubiquitously expressed mimicry transgenes: *MIM159, MIM164*, and *MIM319*. Each of these shows distinct developmental alterations, including hyponastic leaves in *MIM159*, increased leaf serrations in *MIM164*, and reduced fertility in *MIM319* lines (Todesco et al. 2010). We found that the *hws-1* mutation could suppress the characteristic phenotypes of all three mimicry lines (Fig. 2B), implying HWS action upstream of *MIM156*-specific factors like the miR156-targeted *SPL* transcripts. As expected, presence of the *MIM319* and *MIM159* transgenes did not affect the *hws* abscission phenotype (Fig. S2 and see below).

To confirm that HWS acts upstream of miRNA-regulated transcription factors, we introduced a miR156 resistant SPL9 (rSPL9) transgene into *hws* mutants. This transgene expresses a version of SPL9 that avoids regulation by miR156 due to the presence of five base substitutions in its miR156 target site (Wang et al. 2008). Like *MIM156*, rSPL9 plants accumulate higher levels of SPL9 and display *MIM156*-like phenotypes, including spoon-shaped cotyledons and a slower leaf initiation rate (Fig. 2C). We found that the rSPL9 phenotype was similar in wild-type and *hws-5* backgrounds (Fig. 2C), pointing to a role for HWS upstream of miRNA target stability and/or activity.

We speculated that a more general role of HWS could be to impede MIM action at the level of the *IPS1*-based MIM transcript. In this case, MIM transcripts and the endogenous *IPS1* would be affected similarly by *hws* mutation. We thus monitored *IPS1* steady-state levels, both under normal conditions and phosphorus (Pi) starvation, since the latter induces endogenous *IPS1* expression and hence facilitates detection of the otherwise lowly expressed gene (Martín et al. 2000). Independent of Pi supply, *IPS1* accumulation was increased in *hws-5* compared to the wild type (Fig. 2D). Overabundance of mimicry transcripts (i.e., *IPS1*) should lead to more efficient miRNA sequestration, and consequently a release of target suppression. Levels of *PHO2*, the endogenous target of *IPS1*-bound miR399, however, remained unaffected (Fig. 2D). This may indicate a defect in miRNA-mediated target regulation, or result from feedback regulation within the *IPS1-miR399-PHO2* module buffering fluctuations of *PHO2* accumulation (Franco-Zorrilla et al. 2007; Fujii et al. 2005; Bari et al. 2006; Chiou et al. 2006). We therefore turned to analysis of plants harboring the engineered MIM transgenes, which are supposedly uncoupled from endogenous, *IPS1*-promoter-based feedback loops.

The effects of the *hws-5* mutation on engineered *IPS1*-based transcripts was variable (Fig. 2E), but miR156 and miR164 over-accumulated, while expression of their targets was reduced in *hws-5*; *MIM156* and *hws-5; MIM164* plants, respectively, indicating suppression of the MIM phenotype (Fig. 2E).

### HWS reduces miRNA accumulation

The aforementioned observations implied that HWS could be involved more broadly in miRNA regulation, a notion supported by phenotype comparisons. Phenotypic defects in *hws* mutants partially resemble those observed in mutants defective in miRNA biogenesis, or those with altered levels of miR164, a miRNA unrelated to our genetic screen. In *hws* mutants, serrations are reduced (Fig. S3A), a trait that is often affected in miRNA biogenesis mutants (Laubinger et al. 2008; Morel et al. 2002). Moreover, both the ‘skirt’ and cauline leaf fusions to the stem have been described as results of miR164 overexpression (Schwab et al. 2005; Mallory, Reinhart, et al. 2004). Further, when we overexpressed the *HWS* coding sequence from the constitutive CaMV 35S promoter, plants developed severe abnormalities, including upwards-pointing, highly serrated and hyponastic leaves, similar to what was described earlier (Fig. 3A, (Gonzalez-Carranza et al. 2007)). This was also reminiscent of mutants with impaired miRNA activity, for example *hyl1-2, ago1-25* and *ago1-27* or *hasty-3* (*hst-3;* Fig. S3B, (Morel et al. 2002; Vazquez et al. 2004; Bollman et al. 2003)). We therefore decided to test whether miRNAs were directly affected in *HWS*-overexpressing plants as well as *hws* mutants. Using RT-qPCR and small RNA Northern blots, we observed that while miRNA levels tended to be slightly upregulated in *hws* mutants, they were generally downregulated in *35S::HWS* plants compared to the wild-type control (Fig. 3B, D).We also saw a decrease in the levels of miRNA-targeted transcripts of *AGO1* (miR168), *SPL3* (miR156), *TEOSINTE BRANCHED1, CYCLOIDEA and PCF 4* (*TCP4*, miR319), *LEAF CURLING RESPONSIVENESS* (*LCR*, miR394) and *TARGET OF EARLY ACTIVATION TAGGED 2* (*TOE2*, miR172) in *hws* mutants, and a matching increase in *35S::HWS* (Fig. 3C). In addition, *IPS1* accumulation in *35S::HWS* was similar to what was seen in wild type, whereas *PHO2*, the *IPS1*/miR399 target, was upregulated, independent of Pi supply (Fig. 2D). We thus hypothesized that HWS plays a more general role in the miRNA pathway.

**Figure 3.**
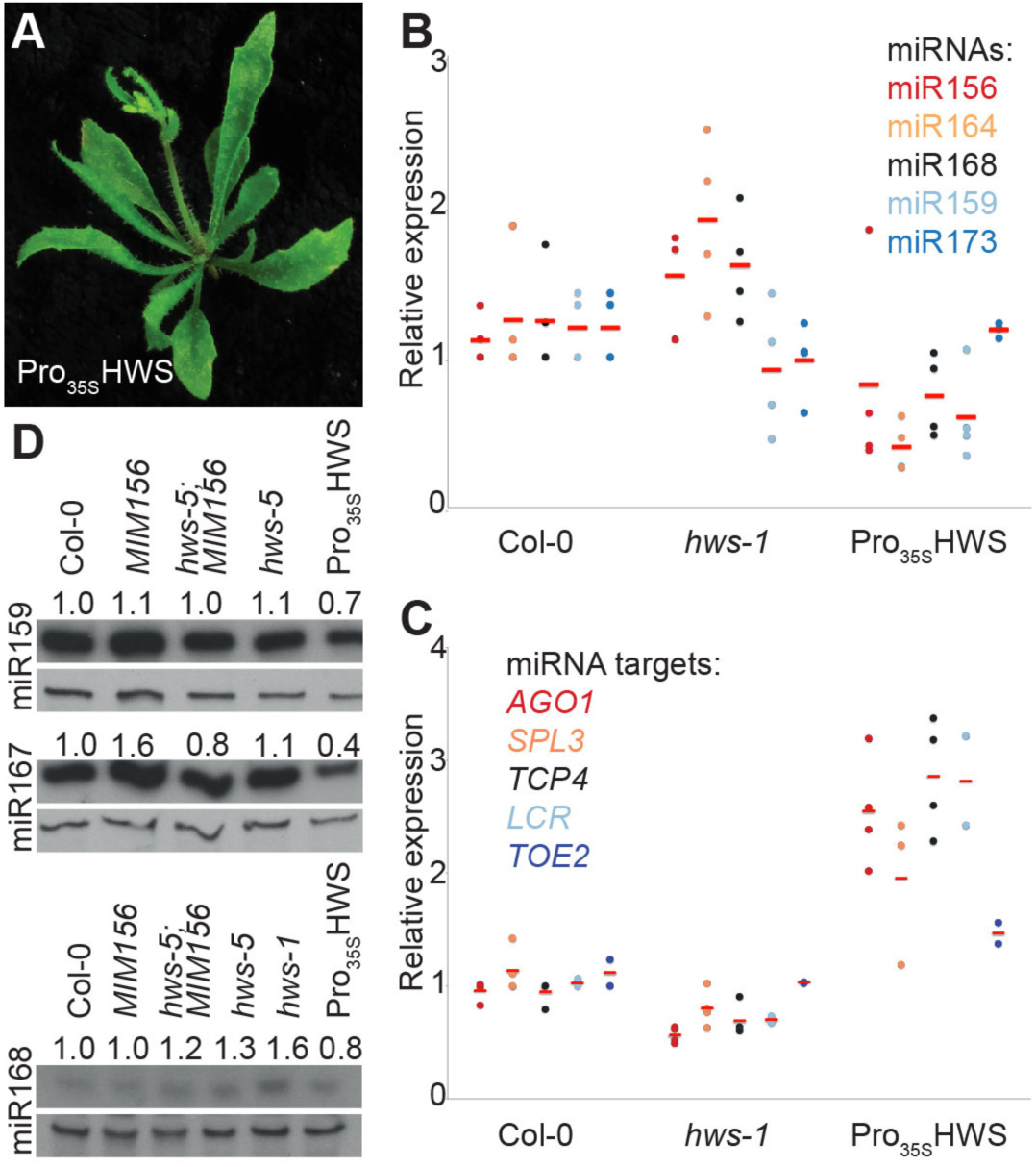
Effects of *hws* and *HWS* overexpression on miRNA and miRNA target steady state levels. (*A*) Phenotype of 28-day old T_1_ *35S::HWS* plant in the *hws-1* background. Note hyponastic, serrated leaves. (*B* and *C*) Levels of mature miRNAs and miRNA targets in Col-0, *hws-1* and *35S::HWS* as measured by RT-qPCR. Dots represent biological replicates (n=3-4), bars indicate mean of biological replicates as means of technical replicates. (*D*) Mature miRNA levels in Col-0, *hws-1, hws-5, MIM156, hws-5; MIM156* and *35S::HWS* as determined by RNA blotting.

The particularly pronounced effects on miR164 and miR156 might be attributed to *HWS* expression being specifically elevated in regions where miR164 and miR156 are expressed. Both miR156 and miR164 accumulate to high levels in emerging leaves (Bazzini et al. 2009; Wang et al. 2009; Wang et al. 2008; Wu & Poethig 2006; Nikovics et al. 2006), with miR164 also showing a more restricted expression pattern around the veins and the points of leaf serration (Fig. S3D, (Nikovics et al. 2006)). A similar expression pattern of the GUS reporter was seen in Pro_*HWS*_::GUS plants (Fig. S3D). This is in agreement with the reduction of serration found in *hws* plants, as in the emerging leaves of wild-type plants, miR164 acts as a suppressor of leaf serration by targeting members of the *CUP-SHAPED COTYLEDON* (*CUC*) family of transcription factors (Fig. S3A, (Nikovics et al. 2006)).

### Genetic interactions with other miRNA factors

To further substantiate a connection between *HWS* and miRNA biogenesis factors, we tested for genetic interactions. For this purpose, we combined the *hws-1* allele with different mutant alleles of *AGO1* (*ago1-25, ago1-27;* (Morel et al. 2002)), *ABA HYPERSENSITIVE1* (*ABH1, abh1-753;* (Laubinger et al. 2008)), *HST* (*hst-3;* (Bollman et al. 2003)), *HYL1* (*hyl1-2;* (Vazquez et al. 2004)), and *SE* (*se-3;* (Grigg et al. 2005)) (Fig. 4). In all cases, doubly-homozygous F_2_ plants resembled the single mutants with defects in the miRNA biogenesis pathway (Fig. 4A). Moreover, the *hws* ‘skirt’ phenotype was reduced (Fig. 4B), suggesting that the miRNA factors are epistatic to HWS.

**Figure 4.**
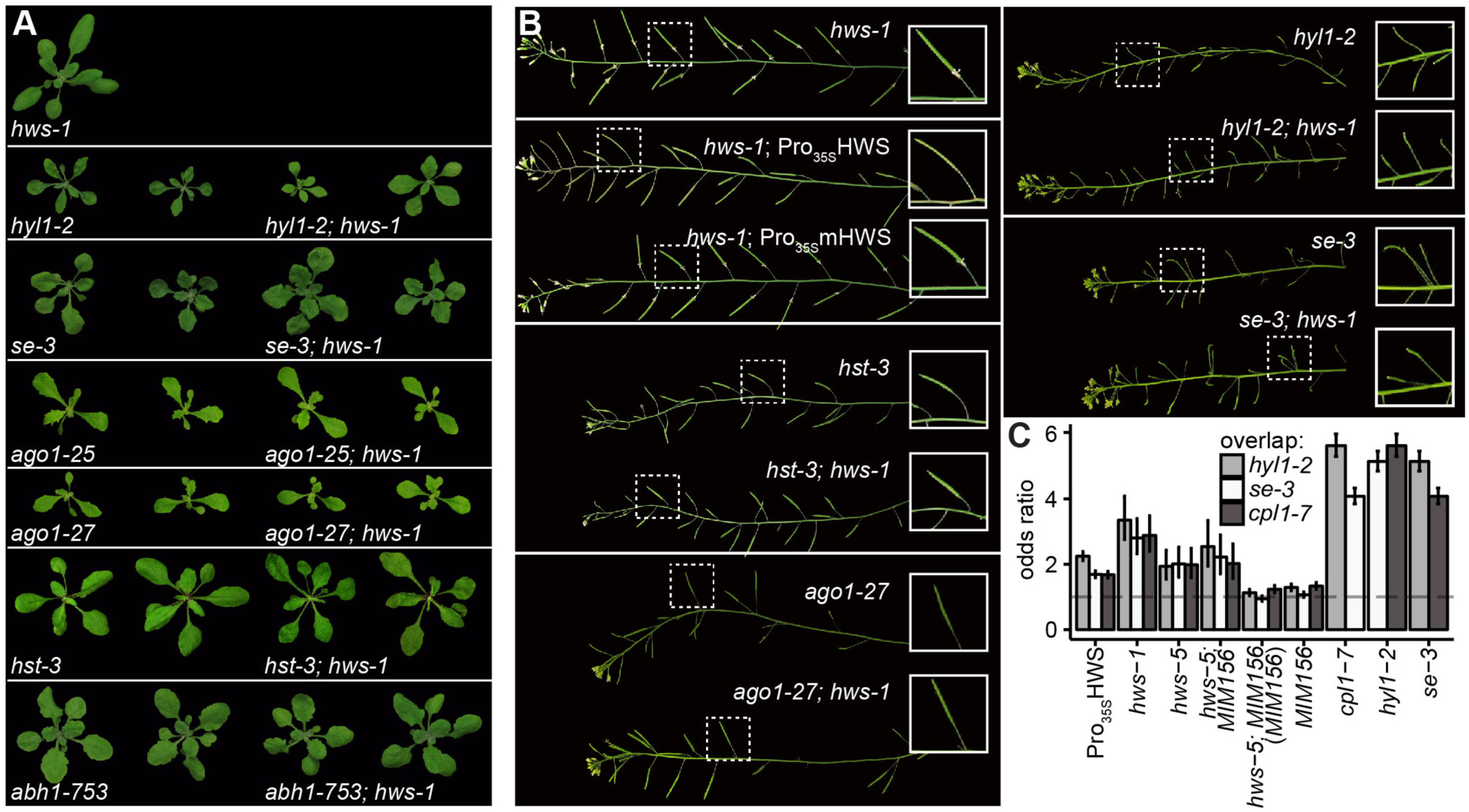
Epistasis analysis of *hws-1* and mutations in major miRNA biogenesis factors. (*A*) Rosette phenotype of single and double homozygous F_3_ plants between *hws-1, hyl1-2, se-3, ago1-25, ago1-27, hst-3* and *abh1-753* at ∼21 DAS. (*B*) Abscission phenotype of double mutants of *hws-1, hws-5, hyl1-2, hst-3, se-3, ago1-27* as well as T_1_ of *35S::HWS* and *35S::mHWS* in *hws-1* background. (*C*) Odds ratio of enrichment of differentially expressed genes also found in the miRNA-mutants *hyl1-2, se-3* and *cpl1-7* as calculated by two-tailed Fisher’s exact test in *35S::HWS, hws-1, hws-5, hws-5; MIM156* (compared to Col-0 and to *MIM156;* n=2), *MIM156* and miRNA-mutants *cpl1-7, hyl1-2* and *se-3* (n=3, data from (Manavella et al. 2012)), with 95% confidence intervals. For p-values, see Table S2.

We reasoned that functional relatedness would be reflected in a substantial overlap of differentially expressed (DE) genes, and thus compared those observed in *hws* to DE genes in the known miRNA mutants *hyl1-2, se-3* and *cpl1-7* (Manavella et al. 2012). We indeed observed that shared misexpression was twice as likely as expected by chance, not too dissimilar from pairwise comparisons among the three previously described mutants (ca. fourfold likelihood of shared misexpression; Fig. 4C; for p-values of two-tailed Fisher’s exact test and odds ratios see Table S2). This trend was also visible in *hws-5* and the suppressor line *hws-5; MIM156* (both compared to Col-0 to identify DE genes), even though the latter has the lowest fraction of DE genes within our samples (Fig. 2A). Conversely, DE genes in *MIM156* did not show this bias, which is not unexpected as only a single miRNA is directly affected by the transgene.–Genes differentially expressed between *hws-5; MIM156* and *MIM156* (i.e. genes affected by *hws* in general, but including those differentially expressed in the *MIM156* background in particular) did not overlap more with those observed in *hyl1-2, se-3* and *cpl1-7* than expected by chance. Nevertheless, these observations support a postulated role of HWS in miRNA biogenesis.

Owing to its F-box domain, HWS’ mode of action could involve an SCF-complex, as HWS has been shown to interact with the common SCF component *Arabidopsis*-Skp protein ASK1 (Ogura et al. 2008; Kuroda et al. 2002). There is precedence for involvement of F-box proteins in miRNA function, examples being the viral silencing suppressor P0 (Pazhouhandeh et al. 2006; Bortolamiol et al. 2007; Baumberger et al. 2007) and F-BOX WITH WD-40 2 (FBW2, (Earley et al. 2010)). Full-length *35S::HWS* complemented the *hws-1* phenotype, whereas transformants expressing a version that lacks the F-box domain (*35S:mHWS* transgene, Fig. 5A) retained the hws-characteristic ‘skirt’ (Fig. S1A, B). Furthermore, *35S::mHWS* did not induce the *hyl1-2* / *ago1*-like leaf phenotypes observed in *35S::HWS* plants (Fig. 3A, 5B). Consistently, steady-state levels of miRNAs and their targets in *35S::mHWS* were more similar to wild type (Fig. S3E, F).

**Figure 5.**
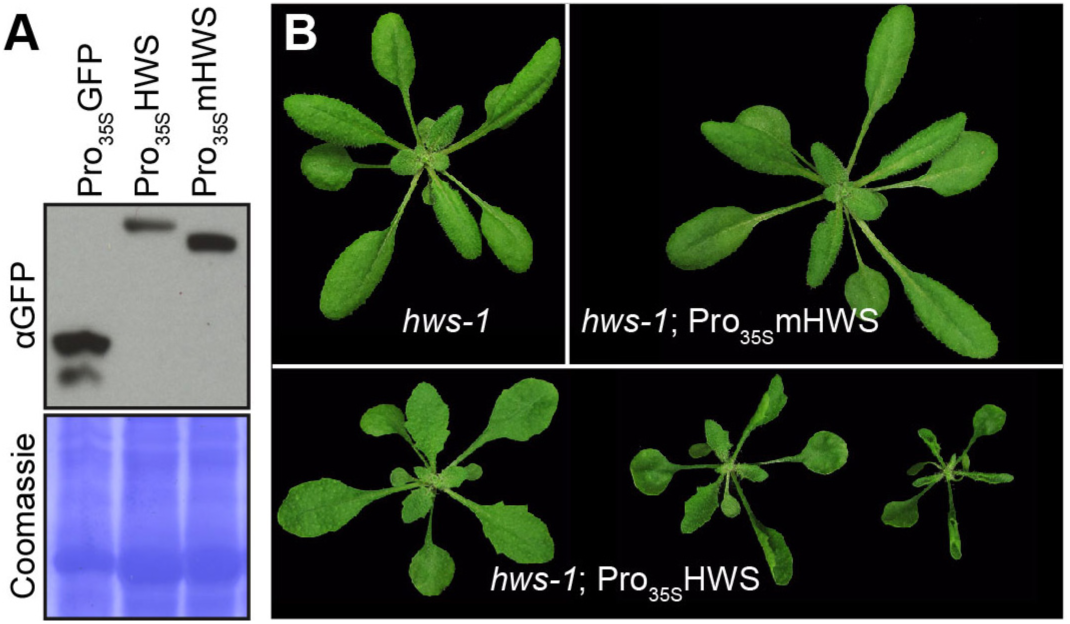
*HWS* overexpression with and without the F-box domain. (*A*) Protein levels of HWS-transgenes in transgenic background from *Arabidopsis* T_1_ and a stable *35S::GFP* line. Coomassie stainings show equal loading. (*B*) Rosette phenotype comparison of *hws-1* and *hws-1* with *35S::HWS* or *35S::mHWS* at ∼21 DAS.

To detect HWS-interactors *in planta*, we harvested rosette leaves from *35S::GFP-HWS* plants, immunoprecipitated the fusion protein with a GFP-antibody and performed mass spectrometry. As controls, we included tissues from both *35S::GFP-mHWS* and *35S::GFP*. Enrichment of ASK1 and two other SCF-complex proteins, ARABIDOPSIS SKP-LIKE 2 (At5g42190) and CULLIN1 (At4g02570), in the *35S::GFP-HWS* fraction, but in neither of the two controls, supported HWS function as a classical F-box protein (Table S3). Beyond this, we did not observe strong associations of HWS with known miRNA-related proteins. Hence, it is possible that HWS contributes to the miRNA pathway through a protein not yet described in this context; that the interaction with an already known factor is rather weak or transient, as described for other F-box proteins (Earley et al. 2010; Coyaud et al. 2015); or that the concentration of the interactor in entire leaves is low and therefore not detectable by our approach. We thus looked for interaction partners in a more direct way, specifically targeting known proteins of the miRNA pathway using a Y2H assay (Manavella et al. 2012; Fields & Song 1989). We could, however, not find a physical interaction between the HWS protein and any of the miRNA biogenesis factors tested (Fig. S4).

One of the miRNA-related proteins detected via MS - albeit at lower levels also in the controls - was AGO1. It is known to associate with several F-box proteins, among them FBW2, which destabilizes it, and the viral suppressor P0 (Earley et al. 2010; Bortolamiol et al. 2007; Baumberger et al. 2007; Pazhouhandeh et al. 2006; Csorba et al. 2015). Similarities between the HWS overexpression phenotype and *ago1* supported a potential functional connection, and AGO1’s role in miRISC assembly fits with presumed HWS action upstream of miRNA targets. However, we did not observe a direct interaction of HWS with either full-length AGO1 (Fig. S4) or the AGO1 N-domain alone (Fig. S5A). This was not entirely unexpected, as F-box proteins often recognize only post-translationally modified targets, and P0 and AGO1 also do not interact with each other in yeast (Bortolamiol et al. 2007; Petroski & Deshaies 2005). We were, however, unable to detect association of HWS and AGO1 by co-immunoprecipitation.

If an interaction of HWS with AGO1 is too labile or transient for robust detection, or indirect, we might still see effects on AGO1 stability, given that they are not masked by the complex feedback loops balancing AGO1 abundance (Vaucheret et al. 2004; Vaucheret et al. 2006). While we observed a positive influence of the presence of HWS’ F-box on HWS stability, AGO1 levels remained unaltered (Fig. 5A, S5B). Ubiquitination by a HWS-associated SCF-complex could, apart from targeting for degradation, affect AGO1 function in other ways, such as altering cellular targeting that was previously implicated in miRISC function (Mukhopadhyay & Riezman 2007; Li & Ye 2008; Brodersen et al. 2012; Gibbings et al. 2009). However, both overall as well as AGO1-specific ubiquitination levels appeared unchanged when HWS was overexpressed (Fig. S5C).

### Widespread HWS action with pronounced effect on the miRNA context

To further pinpoint the function of HWS in the miRNA pathway, we compared transcriptomes of *hws-1* and *hws-5*, the *MIM156* line and the original suppressor line *hws-5; MIM156*, as well as *35S::HWS* and the Col-0 wild type (Fig. 6). In agreement with the morphological phenotypes, the fraction of genes with significant transcript level changes relative to the wild type (adjusted p-value <0.05) was largest in *35S::HWS* and *MIM156* (25% and 15% of all DE genes, respectively), compared to more limited changes in both *hws* mutants (∼2%, Fig. 2A, Table S4).

**Figure 6.**
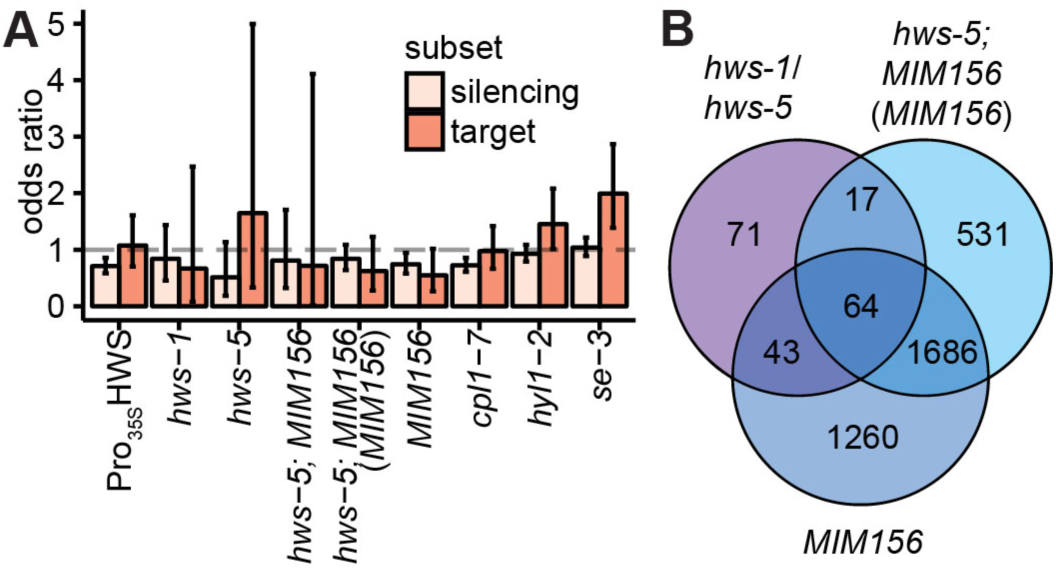
Transcriptome effects of *hws*. (*A*) Odds ratio of enrichment of differentially expressed genes in the silencing or miRNA target gene subset as calculated by two-tailed Fisher’s exact test in *35S::HWS, hws-1, hws-5, hws-5; MIM156* (compared to Col-0 and to *MIM156;* n=2), *MIM156* and miRNA-mutants *cpl1-7, hyl1-2* and *se-3* (n=3, data from (Manavella et al. 2012)), with 95% confidence intervals. For p-values, see Table S2. (*B*) Venn diagram of differentially expressed genes in the *hws* mutants combined compared to *MIM156* and *hws-5; MIM156*.

Looking closer at the DE genes, we classified a set as broadly involved in RNA silencing, comprising also an additional subset of known miRNA targets (Table S6). We used those subsets to determine if genes regulated in a *HWS*-dependent manner would also be indicative of its role as a miRNA factor, and hint at immediately affected processes. We did not observe an over-representation of silencing-related genes among those differentially expressed in *hws* lines - and neither saw such an enrichment in the known miRNA mutants *cpl1-7, hyl1-2*, and *se-3* (Fig. 6A, (Manavella et al. 2012)). The over-representation of miRNA targets was significant in *se-3* and *hyl1-2* (p-value of two-tailed Fisher’s exact test odds ratios p<0.0005 and p<0.05), while *cpl1-7* displayed a similar profile as *35S::HWS*, lacking miRNA target enrichment (Fig. 6A). This suggests that the ‘silencing’ and ‘miRNA target’ annotation we used are not adequate criteria to broadly determine miRNA-relatedness in functionally diverse miRNA mutants. Further, it indicates that the RT-qPCR approach in our case has higher sensitivity than RNA-seq for detecting subtle changes in miRNA target levels.

We therefore turned to categorize transcript profiles of plants with altered *HWS* activity directly, based on transcriptome similarities and dissimilarities in the different backgrounds (Fig. 6B). About half of the DE genes in the two *hws* mutants overlapped and were changed in the same direction (Fig. S6, Table S5). Among those 195 broadly *HWS*-dependent genes, nine were also detected in mass spectrometry (Table S3), and four were found in our ‘silencing’ list (Table S6). Amidst them were the miRNA-function related *GLYCINE-RICH RNA-BINDING PROTEIN 7* (*GRP7*), and *GRP8* (At2G21660 and At4G39260, respectively), and the metabolically active *HOMOCYSTEINE S-METHYLTRANSFERASE1* (*HMT-1*, At3G25900), *GLUTATHIONE S-TRANSFERASE TAU 5* (*GSTU5*, At2G29450) and *GST PHI 4* (*GSTF4*, At1G02950) (Köster et al. 2014; Ranocha et al. 2000; Wagner et al. 2002; van Nocker & Vierstra 1993). The latter two, *GSTU5, GSTF4*, as well as *GRP7*, were also among the 81 differentially expressed genes overlapping in *hws-1, hws-5* and *hws-5; MIM156* (Table S5).

Most expression differences between the suppressor line *hws-5; MIM156* and the *MIM156* parent can be attributed to the phenotypic reversion of *MIM156*-related developmental alterations, and therefore are similarly seen when comparing *MIM156* to the Col-0 wild type (Fig. 2A, 6B). A fraction of DE genes is unrelated, and therefore potentially indicative of HWS downstream effects independent of alterations in miR156 activity, maybe involving other miRNAs (Table S7). Seventeen of those genes are also changed in both *hws* mutants, among them *GRP7* (Table S8).

To find genes regulated by HWS irrespective of the background context, including those responsive to alterations in miR156 (*MIM156* line), we overlaid genes changed in *hws-5; MIM156* with respect to its *MIM156* parent, genes changed in both *hws* mutants (relative to the Col-0 wild-type), and genes changed in the *MIM156* line (also relative to Col-0). The resulting gene list includes *GSTU5, GRP7* and 79 additional genes, of which around half are annotated as enzymes (TAIR, (Berardini et al. 2015); Table S9). Of those, 39 (48%) were also changed in at least two of the previously described miRNA mutants (*hyl1-2, se-3, cpl1-7*). Assuming that at least HYL1 and SE primarily function in (mi)RNA metabolism, the overlap of numerous DE genes, many encoding regulatory proteins, in *hws* and the miRNA mutants could suggest a merging point upstream of those, possibly affecting general metabolic activity.

## Discussion

MiRNA production and function involve a multitude of both general and more specialized factors (Reis et al. 2015; Achkar et al. 2016; Rogers & Chen 2013). Using a *MIM156*-based genetic screen, we have identified the F-box protein HWS as a new factor involved in plant miRNA biology. Both genetic and molecular evidence support a role of HWS in miRNA-dependent processes that goes beyond miR156.

Our observation that *MIM156* levels are reduced in *hws* mutants (Fig. 2E, Fig. S1G) suggests that feedbacks between miR156 and *MIM156* extend beyond this specific context, affecting the equilibrium between MIM and miRNA accumulation. This raises the question where HWS interferes with this balance, and if it is acting down- or upstream of the miR156/SPL regulon. Suppression of additional, miR156-unrelated MIM-phenotypes by *hws* moves HWS away from a highly specialized role in the miR156 context and hints at a more general effect on miRNAs or their targets (Fig. 2B, S2).

Overlap of transcriptome changes between *hws* and *se-3, hyl1-2* and *cpl1-7* – mutants with defects in miRNA biogenesis – (Fig. 4C) further points towards involvement of all factors in a common pathway, possibly upstream of the miRNA-target interaction level. Epistasis of several miRNA related mutations over *hws-1* supports this notion: combination of *hws-1* with such mutants does not change their respective phenotypes, but leads to suppression of the characteristic *hws* ‘skirt’ phenotype (Fig. 4A, B). The ‘skirt’ phenotype is likely caused by overaccumulation of miR164 in floral organs, as it is lost in the presence of *MIM164* (Fig. S2, S3C). This is consistent with earlier observations showing that continuous overexpression of miR164b, as well as *cuc1 cuc2* double mutants, induce fused sepals and stamens and hence floral ‘skirts’ (Mallory, Dugas, et al. 2004; Hibara et al. 2006; Nikovics et al. 2006). It also correlates with the observed positive effect of *hws* on miRNA abundance (Fig 3B, D).

Involvement of HWS in miRNA biogenesis or action and analogous shifts in the MIM/miRNA balance in *hws* could explain the observed suppression of various MIM phenotypes. MIMs largely function as miRNA sponges, which have also been described for animal miRNAs (Ebert et al. 2007), specifically sequestering the miRNAs they can bind to. Since these ‘sponges’ can sequester only a limited amount of miRNAs, overexpression of miRNAs can counter their effects, a scenario potentially reflected in the miRNA-overaccumulating (Fig. 3B, D) and MIM-suppressing *hws* mutant. The effects of MIM constructs on the cognate miRNAs differ substantially: some are greatly reduced, whereas others are only mildly affected (Todesco et al. 2010). Whether this reflects differential efficacies of MIM constructs, or how essential a miRNA is, is not known. Thus, even a small change in MIM transcript levels as in *hws-1; MIM164* (Fig. 2E) could already be sufficient to tip the equilibrium between MIM and miRNA necessary for MIM function, and cause suppression of the charismatic phenotype.

Moreover, consistent with a broader role of HWS, transcriptome changes in *hws* indicate that its effects go beyond merely correcting the aberrant gene expression in *MIM156* plants (Fig. 6B, 4C). Since *HWS* does not encode a transcription factor, the observed changes in the transcriptome are likely cascading effects of its actual, direct effect on one or several proteins. Attempts to pinpoint specific HWS-regulated processes through approaches like GO-term enrichment have however been inconclusive.

Hence, HWS targets and its precise function remain largely enigmatic. Still, as miRNA-like phenotypes and mutant rescue in HWS complementation lines depend on the presence of the HWS F-box domain, it seems highly likely that HWS regulates the stability of a protein important for miRNA biogenesis or action (Fig. S1A, B, 3A, S3B, 5B, E, F). HWS was previously shown to possess F-box ‘activity’ (Takahashi et al. 2004; Kuroda et al. 2002) and confirmed to interact with SCF-complex proteins (this work, Table S3, (Arabidopsis Interactome Mapping Consortium 2011)). Other F-box proteins have been implicated in miRNA function before: AGO1, the core protein of miRISC, is targeted for degradation by the viral suppressor and F-box protein P0, and its levels also decrease upon overexpression of the F-box protein FBW2 (Baumberger et al. 2007; Bortolamiol et al. 2007; Earley et al. 2010; Pazhouhandeh et al. 2006; Csorba et al. 2010).

Because mass spectrometry indicated an enrichment of AGO1 in potential *35S::HWS-GFP* complexes, we performed experiments aimed at identifying changes in AGO1 as a function of HWS activity. While these experiments did not reveal an obvious effect, we cannot exclude AGO1 as a true HWS target yet, as detecting such effects is apparently challenging. As an example, an increase of AGO1 levels in *fbw2* mutants was only detected in genetic backgrounds compromised in miRNA biogenesis, but not in *fbw2* single mutants (Earley et al. 2010). F-box proteins often interact with their targets only transiently, complicating detection of these interactions, especially with heterologous approaches like Y2H (Earley et al. 2010; Bortolamiol et al. 2007). HWS interaction with miRNA biogenesis factors, if present, is likely much weaker and more transient than the HWS F-box connection with SCF complexes, as we could detect clear enrichment of both SKPs and CUL in the mass spectrometry analysis.

Since the F-box domain usually provides the interaction interface towards the other SCF complex components, HWS would interact with its target(s) via the Kelch-domain (Imaizumi et al. 2005; Andrade et al. 2001). Elimination of the F-box should stabilize this interaction (Skaar et al. 2013), an assumption that we could so far not confirm in our experiments (Fig. S5B, data not shown). Most likely, we did not yet find a true HWS target, and/or HWS action requires additional, *Arabidopsis-specific* factors, precluding its function in transient assays. Possibly, a bridging factor is necessary for HWS to indirectly affect miRNA biogenesis, or HWS-mediated targeting depends on prior target modifications, as is for example known for within F-boxes common phosphorylation-dependent recruitment (Skaar et al. 2013). As HWS over-accumulated to higher levels without the F-box (Fig. 5A), and since *35S::HWS*, but not *35S::mHWS* plants, display a strong miRNA-related phenotype, it is also possible that HWS is part of a feedback-loop involving miRNAs. Interaction with or recognition of its target(s) might destabilize HWS, explaining the observed stabilized protein levels in plants lacking the F-box. Alternatively, HWS might to a certain amount self-ubiquitinate, or be degraded together with its target.

The *hws* mutant phenotypes are, apart from the characteristic ‘skirts’, rather subtle. Strong defects are visible only in the overexpressor line or within a sensitized MIM context, while the weak, but pleiotropic effects of HWS deficiency are hardly detectable in an otherwise wild-type background. Effects of *hws* beyond *MIM156* suppression hint at a broader impact, possibly on general metabolic pathways. Genes that are differentially expressed in *hws* mutants and at least in two miRNA mutants tend to be related to cell wall, anisotropic growth and wounding, which is in agreement with HWS affecting cell division activity in the root meristem as well as guard cell growth (Yu et al. 2015; Kim et al. 2016). This rather ‘broad action spectrum’ makes it probably difficult to find specific F-box targets. However, the wide range of effects in multiple pathways fits the role observed here of HWS in miRNA biogenesis and beyond, peak *HWS* expression within the highly developmentally active floral organs, as well as the reported effects on general plant and organ growth (Gonzalez-Carranza et al. 2007; Yu et al. 2015), and quiescent-center independent meristem activity in roots (Kim et al. 2016). Further, it is reminiscent of relatively weak, pleiotropic defects in other miRNA factors like *CPL1* (Manavella et al. 2012; Jeong et al. 2013; Cui et al. 2016) and suggests that HWS may affect growth processes via the miRNA pathway, either through direct involvement in miRNA biogenesis and function, or upstream of miRNA-related processes.

## Material and Methods

### Plant material

*Arabidopsis thaliana* seeds of the Col-0 accession were surface sterilized with 10% bleach, 0.5% SDS and stratified for 2 to 3 days at 4°C. Plants were grown at 23°C either on Murashige Skoog (MS) plates (1/2 MS, 0.8% agar, pH 5.7) or in soil in either short day (8 h light / 16 h dark) or long day conditions (16 h light / 8 h dark) in growth chambers with 65% humidity. A mixture of Cool White and Gro-Lux Wide Spectrum fluorescent lights with a fluence rate of 125–175 μmol m−2 sec−1 was used.

For Pi starvation, plants were germinated on MS plates for 7 days, then shifted to plates with full media lacking Pi ((Conn et al. 2013), 0.8% agar, pH 5.7) and grown for 4 more days. *Nicotiana benthamiana* seeds were surface sterilized and vernalized as described above and grown on soil in long day conditions. Mutant alleles *hyl1-2* (N564863, SALK_064863), *abh1-753* (N516753, SALK_016753), *se-3* (N583196, SALK_083196), *ago1-25, ago1-27, hst-3* (N24278), *miR164a-4* and *hws-1* as well as miRNA mimicry lines *MIM156* (N783223), *MIM159* (N783226), *MIM319* (N783243), *MIM164* (N783232), the Pro_*MIR156c*_::GUS, Pro_*MIR164a*_::GUS and the miR156 overexpressor-line *35S::MIR156B* have been described and were obtained either from the Nottingham Arabidopsis Stock Center (NASC) or from colleagues (Rubio-Somoza et al. 2014; Vazquez et al. 2004; Morel et al. 2002; Todesco et al. 2010; Schwab et al. 2005; Gonzalez-Carranza et al. 2007; Franco-Zorrilla et al. 2007; Bollman et al. 2003; Grigg et al. 2005; Nikovics et al. 2006).

For phenotypic analysis of double mutants between *hws-1* and mutant alleles of *AGO1, HYL1, ABH1, HST* and *SE*, F_2_ or F_3_ plants were selected phenotypically, then genotyped to confirm homozygosity of both mutations. The oligonucleotides used for genotyping can be found in Table S10.

### Transgenes

The *HWS* promoter (2460 bp), genomic (4190 bp) and coding sequences (1236 bp) were PCR-amplified from genomic DNA and cDNA, respectively. They were cloned into pCR8GWTOPO and recombined with ProQuest Two-Hybrid System (Life Technologies) and pGREEN vectors (Hellens et al. 2000). The AGO1 N-domain construct was kindly provided by Peter Brodersen. A detailed list of constructs used in this work can be found in the Supplemental Table S11. All oligonucleotides used to amplify the *HWS* fragments are listed in Table S10.

Transient expression in *Nicotiana benthamiana* after *Agrobacterium*-mediated plant transformation has been described (Yang et al. 2000).

### Mutant screen and segregation of *MIM156* transgene

Plants from a stable miR156 mimicry (*MIM156*) line in Col-0 background were subjected to ethyl methanesulfonate (EMS) treatment as described (Weigel & Glazebrook 2002). M_2_ plants grown in SD conditions were visually inspected for suppression of *MIM156* developmental alterations. Candidate plants were crossed to the Ws-0 accession and genomic DNA of 200-300 pooled F_2_ plants was extracted using a CTAB protocol. Sequencing libraries (Illumina TruSeq DNA Sample Preparation Kit) were 10-plexed (Illumina adapters Set A) per flow-cell lane and sequenced on an Illumina HiSeq 2000 instrument to obtain at least 10-fold genome coverage. SHOREmap was used to identify SNPs and mapping intervals (Schneeberger et al. 2009).

The *MIM156* transgene was removed through outcrossing to the Col-0 accession. Presence or absence of the transgene was deduced from BASTA resistance/sensitivity.

### RNA analysis and sequencing

Total RNA was isolated from pools of about 50 plate-grown seedlings 9 days after sowing using TRIZOL reagent (Life Technologies) and DNase A (Life Technologies) treatment according to manufacturer’s instructions. With RevertAid First Strand cDNA Synthesis Kit (Thermo Scientific), reverse transcription was performed on 1-2 μg of total RNA. Quantitative RT-PCR on *HWS*, mature miRNAs, miRNA-precursors and miRNA targets was executed with Maxima SYBR Green 2X Master Mix (Thermo Fisher Scientific) on a CFX384 Real-Time PCR system (Bio-Rad), performing technical triplicates on each sample of biological triplicates using *ACTIN2* (At3G18780) as reference gene. Biological replicates are averaged from technical triplicates, horizontal bars show the mean of the biological triplicates. All oligonucleotides used for RT-PCR experiments are listed in Table S10.

For RNA-seq, total RNA was extracted from pooled flowers, using TRIZOL reagent (Life Technologies) and DNase A (Life Technologies) treatment according to manufacturer’s instructions and a final cleanup using RNeasy Mini spin columns from the RNeasy Plant Mini Kit (Qiagen). Transcriptome libraries were prepared from 1_ug total RNA with a TruSeq RNA Library Prep Kit v2 (Illumina) and single-end sequenced on a HiSeq 3000 with 100 bp reads. Reads were mapped to the TAIR10 *Arabidopsis thaliana* genome and gene expression was quantified using RSEM (v1.2.30, (Berardini et al. 2015; Li & Dewey 2011)). All data analysis after initial read cleanup was executed in R. After exclusion of a contaminated sample, differential expression and all subsequent analysis was conducted with two biological replicates, using DESeq2 (v1.14.1). Differential expression was assessed in comparison to the common wild-type Col-0, and for the suppressor line additionally to *MIM156*. Published transcriptome data from *cpl1-7, hyl1-2* and *se-3* seedlings were used for further comparisons (Manavella et al. 2012).

### Histochemistry

Seedlings from at least five independent GUS reporter T_2_ lines were inspected 10 days after sowing (DAS). Activity of the GUS reporter was assessed as described (Weigel & Glazebrook 2002), using 20 mM potassium-ferro- and 20 mM potassium-ferricyanide.

### Protein analyses

T_1_ seedlings expressing *35S::GFP-HWS* and *35S::GFP-mHWS* were BASTA-selected on soil and harvested at 21 days for total protein extraction from three to six whole rosettes as tissue pools. Protein was extracted from ∼300 to 1000 mg of ground tissue using equal amounts [w/v] of extraction buffer (50 mM Tris pH 7.5; 150 mM NaCl; 1 mM EDTA; 10% [v/v] glycerol; 1 mM DTT; one tablet of Complete Protease Inhibitor Cocktail per 10 ml buffer). Protein concentration was measured using Bradford solution (Bio-Rad). Expression of the fusion protein was tested by Western blot using GFP-trap (ChromoTek), and appropriate pools were chosen for co-immunoprecipitation with GFP-trap or anti-AGO1 (Agrisera) antibodies.

For protein expression analyses, leaves of *Nicotiana benthamiana* transiently cotransformed with *35S:GFP-HWS* or *35S:GFP-mHWS* and *35S:AGO1-HA* were harvested three days after infiltration. Protein abundance was measured by Western blot using anti-AGO1 (1:10000, Agrisera AS09 527), anti-GFP (1:10000, Santa Cruz Biotech, sc-8334) or anti-UBQ (1:2000, Santa Cruz Biotech, sc-8017), equal loading was confirmed using protein staining with either Ponceau red or Coomassie blue.

Interaction experiments in yeast were performed using the ProQuest Two-Hybrid System (Life Technologies) and yeast strain AH109. To reduce autoactivation of some constructs, 5 mM of 3-AT (3-amino-1,2,4-triazole) were added to the selection medium.

### Mass spectrometry

Pools of BASTA-selected T_1_ seedlings expressing *35S::GFP-HWS* and *35S::GFP-mHWS* and GFP-overexpressing control plants were frozen in liquid nitrogen and total protein was extracted from up to 1 g finely ground tissue with equal amounts [w/v] of extraction buffer (140 mM NaCl; 8 mM Na_2_HPO_4_*7H_2_O; 2 mM KH_2_PO_4_, pH 7.4; 1 mM EDTA; 0.1% [v/v] Triton X-100; 1 tablet of Complete Protease Inhibitor Cocktail (Roche) per 10 ml buffer). Protein concentration was measured using Bradford solution (Bio-Rad) and GFP-expression was verified by Western blot.

Total protein extracts were purified using GFP-trap metal beads (ChromoTek). A small fraction was resolved on a PAGE gel for staining with the SilverQuest^™^ Silver Stain Kit (Life Technologies). LC-MS/MS analysis (120 min, Top15HCD) was performed after tryptic in gel digestion, using a Proxeon Easy-nLC (Proxeon Biosystems) coupled to an LTQ Orbitrap Elite mass spectrometer (Thermo Fisher Scientific) (Borchert et al. 2010). Resulting data was analyzed with MaxQuant v.1.2.2.9 (Cox & Mann 2008; Cox et al. 2011). Spectra were searched against an *Arabidopsis thaliana* database including the protein sequences of the HWS::GFP fusion proteins. Raw data were processed with a setting of 1% for the false discovery rate (FDR; Table S3).

## Acknowledgements

We thank Diep Tran for support and suggestions with protein work, Jim Carrington and coworkers for *ago1-25* and *ago1-27*, Jeremy A. Roberts for *hws-1*, Jia-Wei Wang for Pro_*MIR156c*_::GUS and EMS-mutagenized *MIM156* populations, and Patrick Laufs for Pro_*MIR164a*_::GUS seeds, Pablo A. Manavella and Delfina Ré for parts of the ‘silencing gene’ list and Peter Brodersen for the *AGO1* N-domain Y2H construct. We are grateful to Ignacio Rubio Somoza, Pablo Manavella and Moisés Exposito Alonso for valuable discussion, input for experiments, analysis and reviewing the manuscript.

## Competing interests

The authors declare no competing or financial interests.

## Author contributions

M.D.C. and P.L. designed and performed most of the experiments. M.D.C. designed and executed the screen and identified the suppressor. J.H. mapped the suppressor. A.-L. V.d.W. processed transcriptome reads for differential expression analysis, P.L. analyzed transcriptome data. E.D. performed phosphate starvation assays. D.W. supervised research. P.L., M.D.C., R.S. and D.W. analyzed results and wrote the manuscript.

## Funding

This work was supported by the DFG through SFB1101, a DAAD scholarship to E.D., and the Max Planck Society.

## Data availability

RNA-seq reads are available at XXX.

